# Field testing of biohybrid robotic jellyfish to demonstrate enhanced swimming speeds

**DOI:** 10.1101/2020.09.24.312322

**Authors:** Nicole W. Xu, James P. Townsend, John H. Costello, Sean P. Colin, Bradford J. Gemmell, John O. Dabiri

## Abstract

Biohybrid robotic designs incorporating live animals and self-contained microelectronic systems can leverage the animals’ own metabolism to reduce power constraints and act as natural chassis and actuators with damage tolerance. Previous work established that biohybrid robotic jellyfish can exhibit enhanced speeds up to 2.8 times their baseline behavior in laboratory environments. However, it remains unknown if the results could be applied in natural, dynamic ocean environments and what factors can contribute to large animal variability. Deploying this system in the coastal waters of Massachusetts, we validate and extend prior laboratory work by demonstrating increases in jellyfish swimming speeds up to 2.3 times greater than their baseline, with absolute swimming speeds up to 6.6 ± 0.3 cm s^-1^. These experimental swimming speeds are predicted using a hydrodynamic model with morphological and time-dependent input parameters obtained from field experiment videos. The theoretical model can provide a basis to choose specific jellyfish with desirable traits to maximize enhancements from robotic manipulation. With future work to increase maneuverability and incorporate sensors, biohybrid robotic jellyfish can potentially be used track environmental changes in applications for ocean monitoring.

## 1. INTRODUCTION

With ocean acidification altering animal behavior and function [1,2] and temperature-induced biodiversity changes in marine environments [3,4], new tools can expand efforts to track markers of climate change in more sensitive or previously unexplored areas of the ocean [5]. Traditional ocean monitoring tools, such as autonomous underwater vehicles (AUVs) and remotely operated vehicles (ROVs), offer invaluable opportunities to explore the ocean. For example, prior work using AUVs have yielded observations of deep-sea animal communities over multiple decades [6], and ROVs have been used to monitor anthropogenic disturbances of ecosystems [7] and capture gelatinous midwater animals with soft robotic arms [8]. Despite advantages such as speed and reliability [9,10], AUVs and ROVs are still limited in confined spaces and fragile environments, such as near coral reefs or in caves, where debris can cause severe damage to the vehicles [8,11]. These technologies can also cost thousands of dollars and require specialized operational personnel [12].

In conjunction with AUVs and ROVs, other tools that offer alternative strategies can be developed to expand human capabilities to monitor a variety of ocean environments. One potential solution is to take inspiration from biological organisms, which offer advantages in energy efficiency, maneuverability, and stealth compared to extant robotic systems [13,14]. Bioinspired soft robots can potentially address issues in power consumption [13,15] and leave wakes that mimic the wakes of marine life, with potential to minimally perturb surrounding wildlife. Examples of bioinspired aquatic robots include robotic fish [16-18], manta rays [19-21], sea stars [22,23], and jellyfish [24-29], including systems that have been deployed in real-world environments [16,24].

In particular, moon jellyfish (*Aurelia aurita*) are a compelling model organism for building robots because of the limited energy required for locomotion. *A. aurita* is a species of moon jellyfish that comprise a flexible oblate bell, composed of mesoglea (gelatinous structural tissue that primarily comprises water and extracellular proteins) with a singular muscle layer oriented circumferentially on the subumbrellar surface. The animal has eight natural swim pacemakers located on the bell margin, each of which can independently activate to excite the swim muscle. This initiates the power stroke, in which the muscle contracts to decrease the subumbrellar volume and generate thrust to travel forward. The muscle then rests during the relaxation stroke, returning the jellyfish bell back to its relaxed shape [30]. Induced flow from stopping vortices during a relaxed phase of the swimming cycle provides additional thrust at no increased metabolic input. This process, known as passive energy recapture, allows jellyfish to have the lowest cost of transport (COT), defined as the mass-specific energy input per distance traveled, compared to other animals [31].

However, bioinspired robotic constructs mimicking jellyfish still exhibit higher energy costs than their biological analogs [24,26]. An alternative approach is to incorporate live animals into a biohybrid robotic construct, which can then use an inexpensive and simpler microelectronic system to power electrodes that excite an existing biological system, instead of the energy costs and design considerations for using mechanical actuators and chassis. Biological components can also improve damage tolerance using natural tissue regeneration. Instead of relying on the animals’ natural pacemaker system to activate muscle contractions, a robotic system with electrodes that generate square pulse waves of 3.7 V was previously described to incite jellyfish muscle contractions [32]. Prior work has shown that even without arresting endogenous pulses in the animals, driving the jellyfish at various frequencies with the portable swim controller resulted in increased swimming speeds [32].

Although biohybrid designs incorporating live animals are limited by biological constraints, this system can also improve upon biological performance. For example, previous work has demonstrated that by driving jellyfish at faster frequencies than they would normally exhibit themselves, biohybrid robotic jellyfish can increase swimming speeds up to 2.8 times, at only a 10 mW input to the robotic system and twofold increase in metabolic cost to the animal. Biohybrid robotic jellyfish also use less external power per mass than other reported swimming robotics in literature [32]. The ubiquity of jellyfish found at various depths, including thousands of meters below surface level [33], offers opportunities to incorporate biohybrid robots to explore new areas of the ocean in the future. This would require only a hardened microelectronic system, as opposed to an entire robot that could be easily damaged in real conditions.

However, previous demonstrations of biohybrid robotic jellyfish were limited to controlled laboratory experiments. Open questions include how natural environmental conditions, such as current and turbulence, affect swimming performance relative to laboratory results in quiescent conditions, and the feasibility of future ocean monitoring using this integrated swim controller and live animal design. We conducted a series of vertical swimming experiments in the coastal waters of Massachusetts to test the effect of externally driven swim controller frequencies on vertical swimming speeds in the ocean. We hypothesized that even in the presence of surface currents, increasing swimming frequency would increase swimming speeds up until a limit, in which altered swim kinematics would result in decreased swimming speeds.

## 2. METHODS

### (a) Animal care

*A. aurita*, originally housed in facilities at Stanford University (animal husbandry details described in [32]), were shipped overnight to the Marine Biological Laboratory (MBL) in Woods Hole, MA. Animals were subsequently stored at room temperature, 21°C, in standard 5-gal plastic buckets filled with filtered natural seawater from the Atlantic Ocean, and fed naupliar brine shrimp for one hour before daily water changes.

### (b) Biohybrid robotic system

We adapted the robotic system described in [32] for use in field experiments. The swim controller comprised a mini processor (TinyLily, TinyCircuits, Akron, OH, USA) and 10-mAh litihium polymer cell (GM201212, PowerStream Technology Inc., Orem, UT, USA) in plastic housing made entirely from polypropylene pieces (Fig. 1B) sealed with hot melt adhesives, as opposed to the previous design with Parafilm M. The housing was ballasted with stainless steel washers to keep the system neutrally buoyant in seawater. Two electrodes were assembled using perfluoroalkoxy-coated silver wires and platinum rod tips (A-M Systems, Sequim, WA, USA) connected in series to red LEDs (TinyLily 0402, TinyCircuits, Akron, OH, USA) as a visualization tool. Platinum wire tips were hooked to improve attachment (Fig. 1A), an additional design feature to secure the swim controller to the animal in field conditions. Examples of the swim controllers are shown in Fig. 1C.

**Figure 1.**
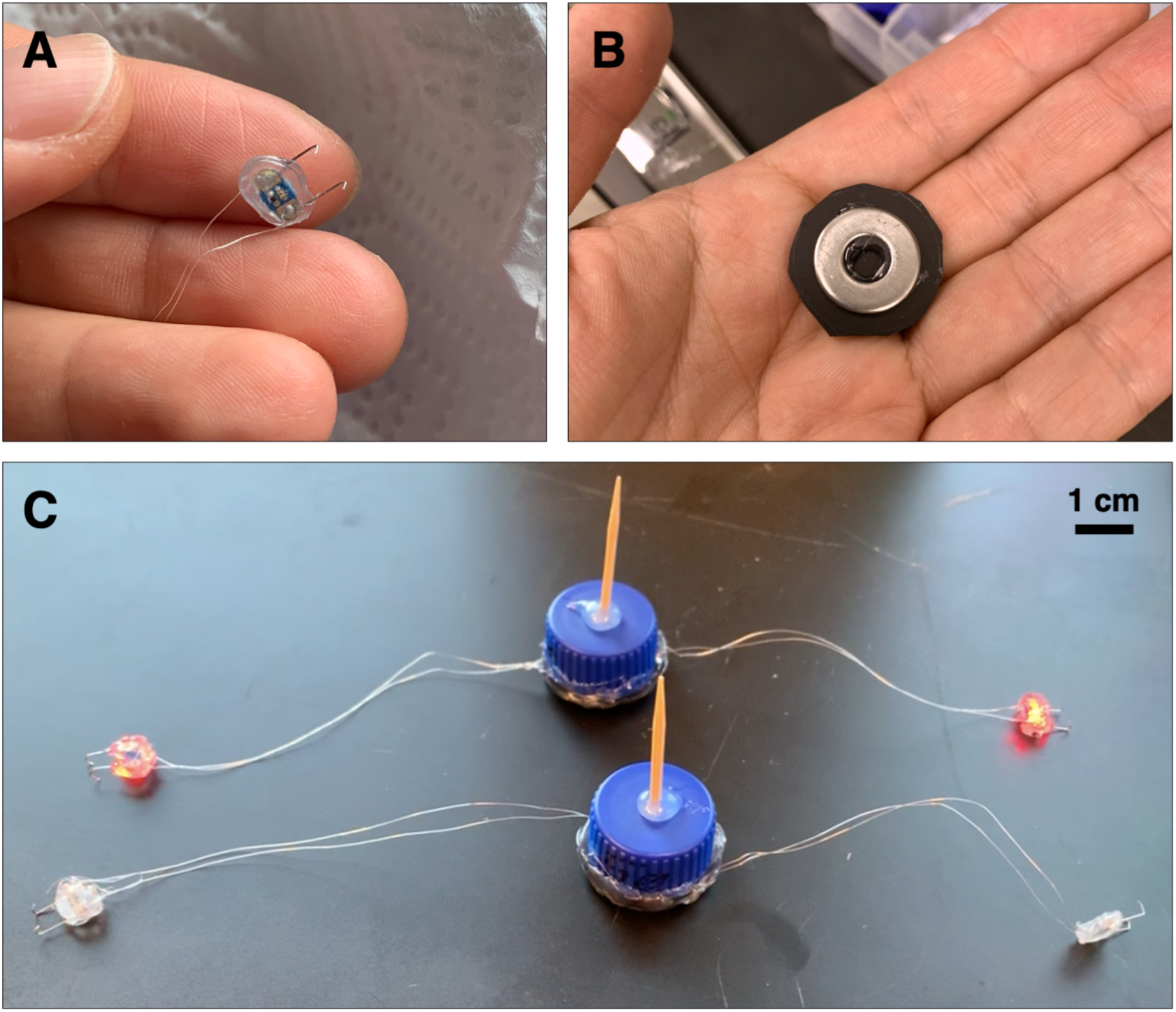
Robotic system using microelectronics. The biohybrid robotic system, adapted from [32], with two new features: (A) hooked electrode wire tips instead of straight tips in the previous design, and (B) a weighted polypropylene cap to change the ballast and improve the robustness of the system under field conditions, as opposed to no flow in laboratory tank experiments. (C) Two of the fully integrated robotic systems with new modifications are shown. The electrodes shown in the back are active (red LEDs are on).

The robotic system was attached to the jellyfish bell in three locations: a wooden pin connected to the housing was inserted into the center of the manubrium from the subumbrellar surface, and each electrode was inserted into the subumbrellar tissue.

### (c) Field experiments

Preliminary field tests were conducted in water < 1 m in depth to determine the appropriate ballast of the system and robustness of waterproofing techniques on active microelectronics. Subsequent field experiments were conducted in 1.6-m depth water in Woods Hole, MA (dive coordinates 41°31’29.1”N latitude, 70°40’23.8”W longitude) and involved a minimum of two scientific scuba divers and one person on shore. One diver maneuvered animals (*N =* 2) into the starting position (initially at the ocean bottom) near a rope with alternating red and yellow markers at every 30.5 cm, as a known scale for image analysis. Another diver operated a camera system to track the animal and the background rope markers as the biohybrid robotic jellyfish swam upwards to the ocean surface. Videos were recorded in 1920×1080 resolution at 30 fps using a Sony AX100 in a Gates AX100 Underwater Housing (Sony, Tokyo, Japan) on the first dive. An additional *N =* 2 animals were recorded on a second dive for further observations of animal behavior, and were recorded on an iPhone XS (Apple, Cupertino, CA, USA) in a Kraken Universal Smart Housing (Kraken Sports, London, Ontario, Canada). A simplified schematic of the experimental setup is illustrated in Figure 2. Animals were monitored to ensure recovery after experiments (for more information, see “Ethical considerations” in Supplementary Material).

**Figure 2.**
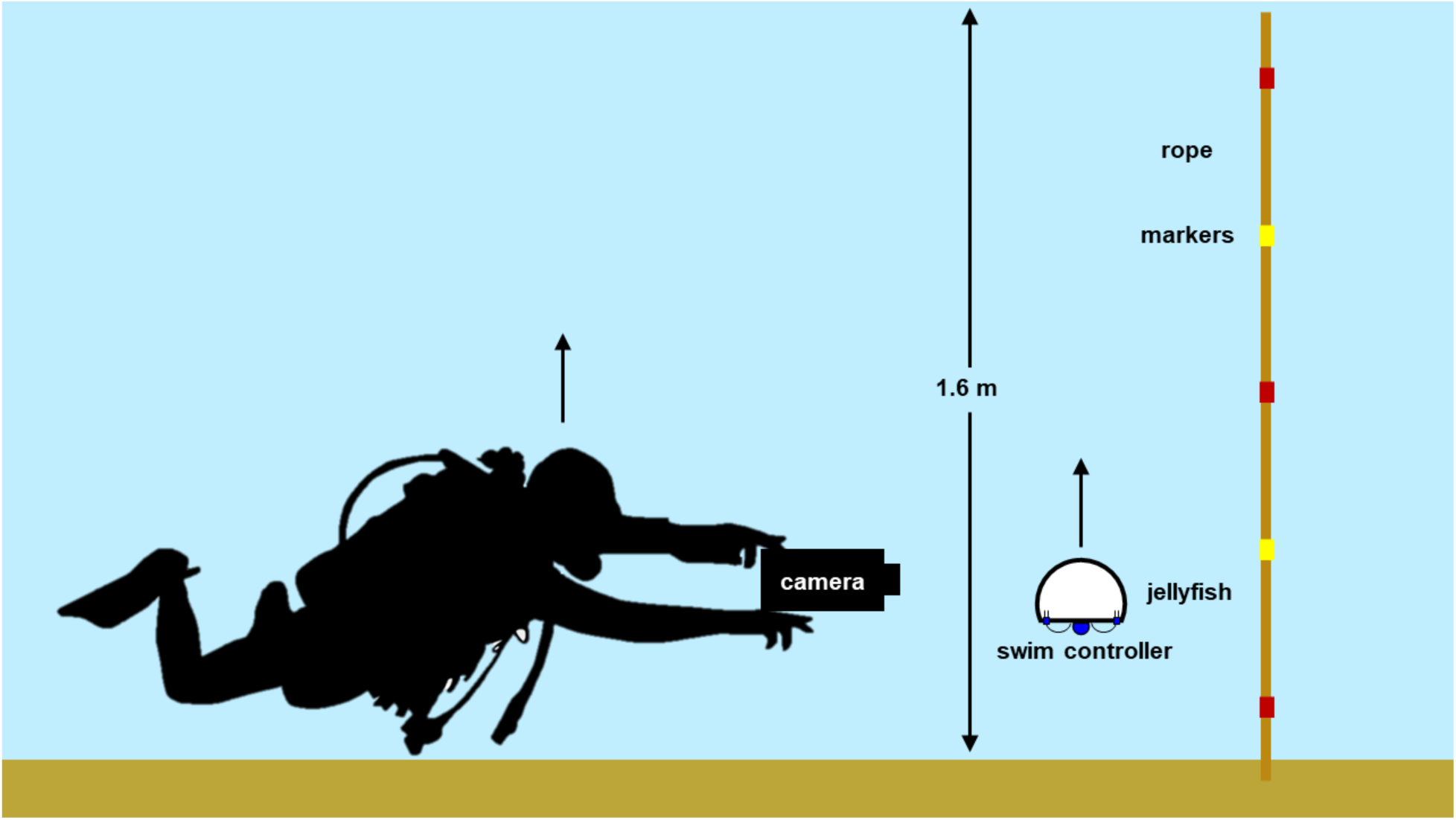
Setup of field experiments. Simplified schematic of the experimental setup, including a scientific diver holding a camera that tracks a biohybrid robotic jellyfish (swim controller and jellyfish) swimming upwards from the ocean floor to the surface, 1.6 m in depth. A rope with alternating red and yellow markers is used to track displacement during image analysis.

Control cases (0 Hz) for each external swim controller frequency (0.50, 0.75, and 1.00 Hz) were tested by cutting both electrode wires, while keeping the electrodes embedded into the animals to maintain neutral buoyancy. In addition to swim controller frequencies (0, 0.50, 0.75, and 1.00 Hz), measured frequencies of the biohybrid robotic jellyfish were determined by counting the number of animals’ pulses within a given time frame. Swim controller frequencies for each animal and wind conditions for Falmouth, MA [34] are listed in Table 1.

**Table 1.** Field test variables. Animals (*N =* 4) and swim controller frequencies tested *in situ*. Bolded conditions were tested on the first day.

| *Jellyfish* | *Swim Controller Frequencies (Hz) Tested* | *Dive Condition Mean Wind Speeds ($m s^{-1}$) and Directions [34]* |
| --- | --- | --- |
| **1** | **0, 0.50, 0.75** | **3.9 $m s^{-1}$**<br>**West-southwest** |
| **2** | **0, 0.50, 0.75** | **3.9 $m s^{-1}$**<br>**West-southwest** |
| 3 | 0, 0.50, 1.00 | 4.6 $m s^{-1}$<br>West-southwest |
| 4 | 0, 0.75 | 4.6 $m s^{-1}$<br>West-southwest |

### (d) Data analysis

Representative images from the videos collected are shown in Figure 3 at various depths during the vertical swimming of each animal. For *N =* 2 animals on the first dive, we tracked centroids of the red and yellow rope markers (see Supplementary Material, Fig. S1A) and housing of the swim controller system (see Supplementary Material, Fig. S1B), assuming pixel-level accuracy in centroids. Vertical displacements of the biohybrid robotic jellyfish over time (Fig. 4) were calculated by determining the position of the biohybrid robotic jellyfish with respect to the rope markers. Vertical speeds were calculated using the vertical positions between subsequent time steps, and averaged to obtain mean vertical speeds at each test condition. Using vertical speeds, enhancement values were calculated as the measured swimming speed at each experimental condition normalized by the baseline swimming speed. The baseline is defined as the swimming speed of the individual biohybrid robotic jellyfish at 0 Hz, the control case.

**Figure 3.**
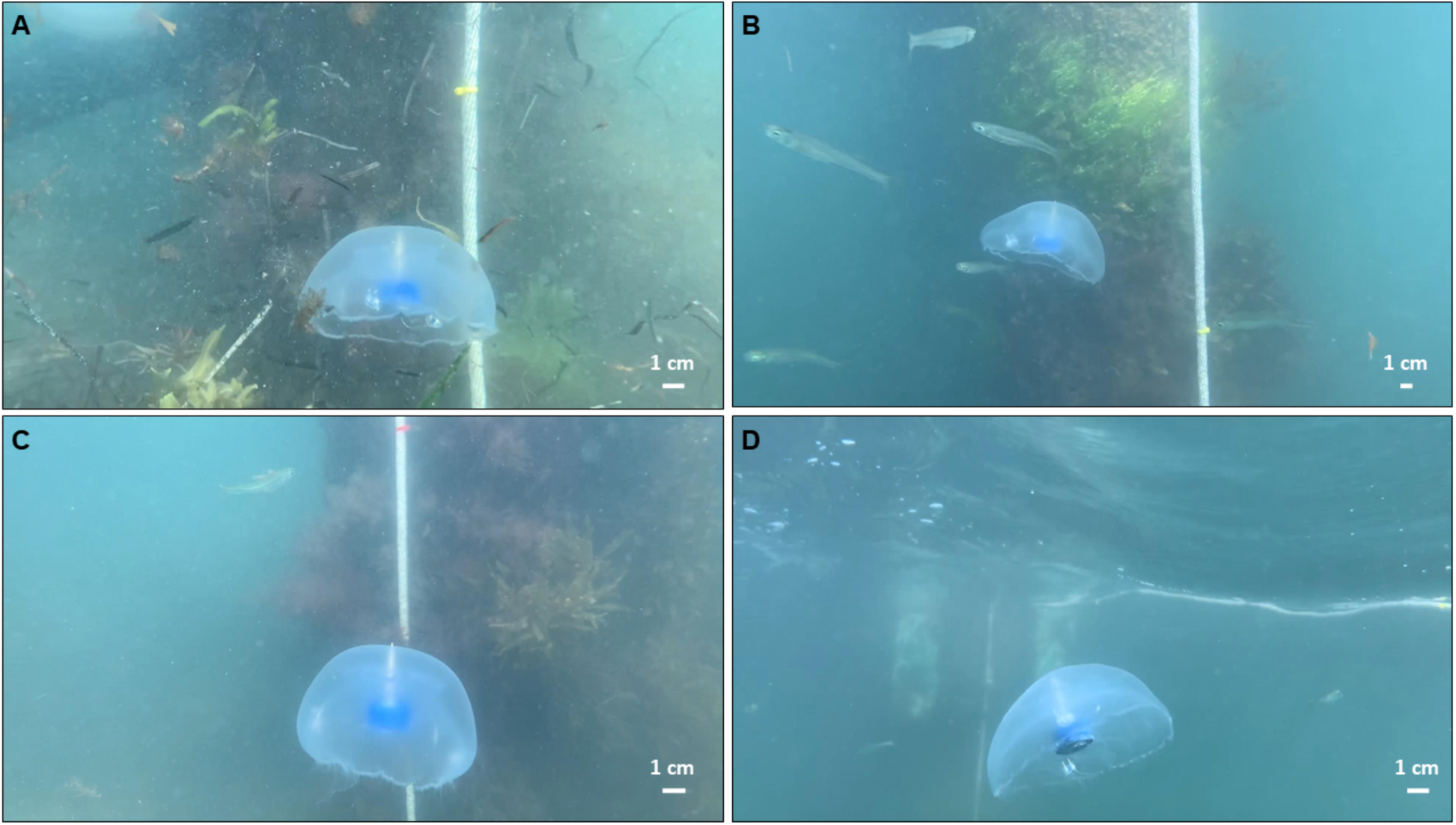
Representative images of biohybrid robotic jellyfish during field experiments. Examples of one biohybrid robotic jellyfish (animal 1) (A) initiated at the ocean bed and stimulated at 0.50 Hz, (B) swimming toward the ocean surface at 0.50 Hz, (C) swimming toward the ocean surface with an inactive robotic system (0 Hz, control), and (D) toward the ocean surface with an inactive robotic system (0 Hz, control).

**Figure 4.**
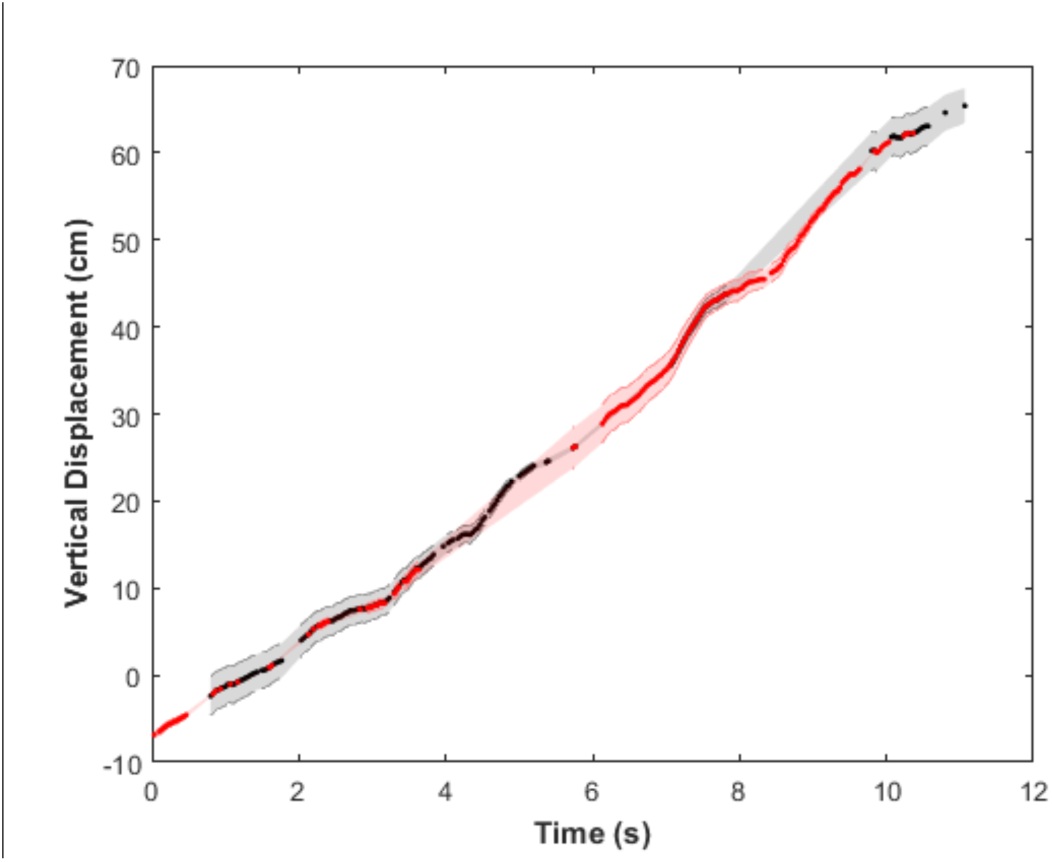
Representative plot tracking animal displacements over time to calculate vertical swimming speeds. Vertical displacement over time of one example jellyfish (animal 1 driven at0.75 Hz) with respect to the rope markers, with the error propagated from conversions in pixel space to centimeter space. Tracks were assembled by stitching vertical positions using both red and yellow rope markers (shown in red and black, respectively, for improved visualization), to show accuracy in overlap.

Similarly, 2D displacement over time was calculated using both vertical and horizontal components (as shown in Fig. 5). Although *N =* 4 animals in total were used over two days, rope markers were not visible in videos of *N =* 2 animals on the second day. However, videos from the second dive provide observational data regarding kinematics and confirm swimming speed estimates.

**Figure 5.**
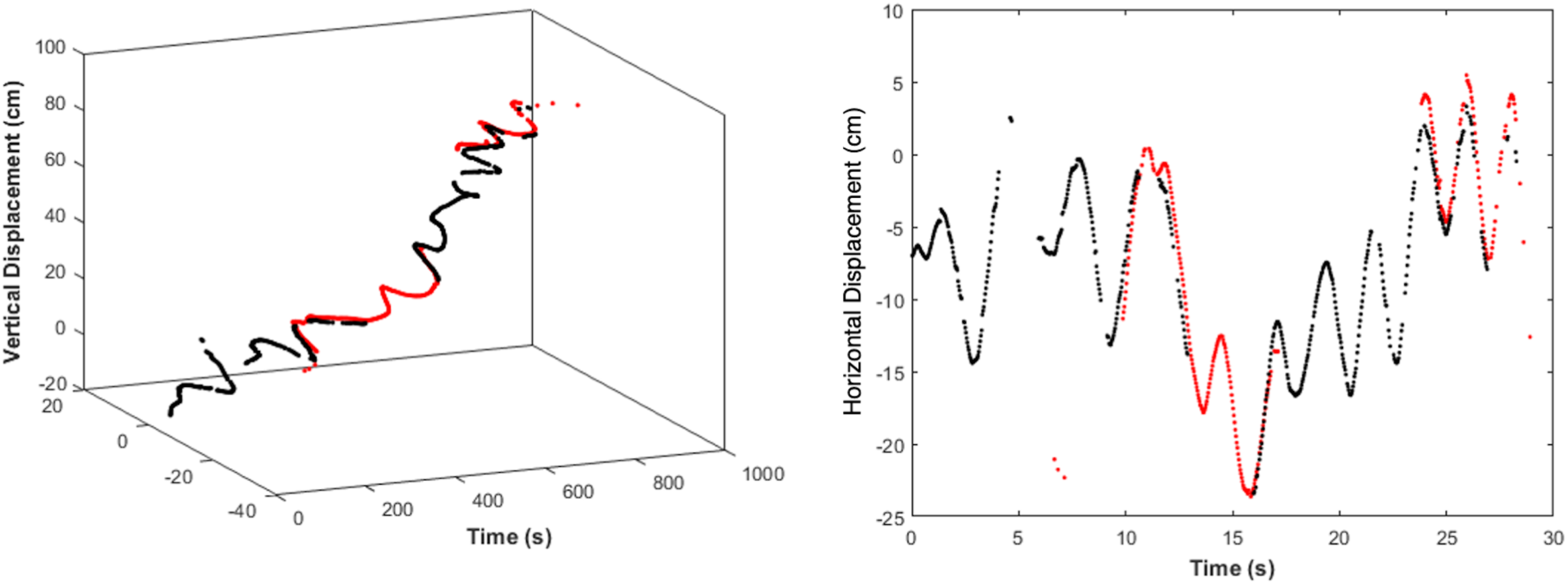
Representative plots tracking animal displacements over time to show horizontal displacements caused by ocean currents. (A) An example 2D displacement over time and (B) the horizontal component over time from one video (animal 1, driven at 0.50 Hz) to show oscillatory effects from primarily horizontal surface currents.

### (e) Hydrodynamic model

As described in [32], a hydrodynamic model was adapted from [35,36] to calculate the velocity (*u*) from a momentum balance using thrust (T), drag (D), acceleration reaction (AR), and inertial forces at a Reynolds number of 325:

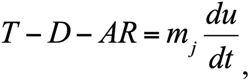

in which

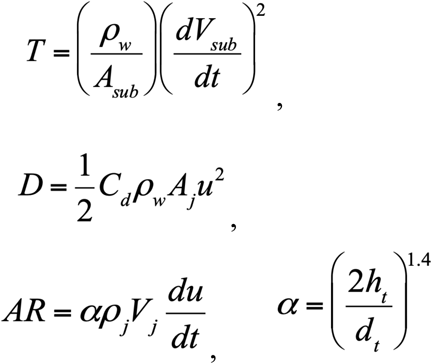

with the following terms:

*m*_*j*_ mass of the jellyfish

*ρ*_*w*_ density of saltwater *=* 1.024 g/cm^3^ at 35 ppt and 21°C

*A*_*sub*_ area of the jellyfish subumbrella

*V*_*sub*_ volume of the jellyfish subumbrella

*C*_*d*_ drag coefficient *=* 0.42

*A*_*j*_ area of the jellyfish

*h*_*t*_ height of the jellyfish

*d*_*t*_ diameter of the jellyfish

*ρ*_*j*_ density of the jellyfish

*V*_*j*_ volume of the jellyfish

Model inputs included both morphological and time-dependent parameters: relaxed bell height (*h*_*r*_) and diameter (*d*_*r*_), maximum change in height between contraction and relaxation states (Δ*h*), maximum change in diameter between relaxation and contraction states (Δ*d*), manubrium tissue height (*h*_*j*_), contraction time (*t*_*c*_) defined as the transition from a relaxed to a contracted state, and relaxation time (*t*_*r*_) defined as the transition from a contracted to a relaxed state.

Velocities from the mechanistic model were calculated for *N =* 2 jellyfish with geometric inputs estimated from experimental videos from the highest measured swimming speeds using ImageJ (National Institutes of Health and the Laboratory for Optical and Computational Instrumentation) and MATLAB (Mathworks). Inputs for the model are listed in Table 2. Mean speeds were calculated from velocities at each time step (30 time steps per second, as a fair comparison to 30 fps in experimental data) for 10 periods.

**Table 2.** Input parameters for the hydrodynamic model. Parameters include measured swimming frequency (*f*), relaxed bell diameter (*d*_*r*_), maximum change in diameter between relaxation and contraction states (Δ*d*), relaxed bell height (*h*_*r*_), maximum change in height between contraction and relaxation states (Δ*h*), manubrium tissue height (*h*_*j*_), contraction time (*t*_*c*_), and relaxation time (*t*_*r*_).

| *Animal* | *f (Hz)* | *d<sub>r</sub> (cm)* | *Δd (cm)* | *h<sub>r</sub> (cm)* | *Δh (cm)* | *h<sub>j</sub> (cm)* | *t<sub>c</sub> (s)* | *t<sub>r</sub> (s)* |
| --- | --- | --- | --- | --- | --- | --- | --- | --- |
| 1 | 0.09<br>0.20<br>0.47<br>0.53<br>0.81 | 11.3 | 6.0 | 4.4 | 1.0 | 2.0 | 0.70 | 0.73 |
| 2 | 0.40<br>0.50<br>0.53<br>0.75 | 9.8 | 5.0 | 4.2 | 1.0 | 2.0 | 0.87 | 0.90 |

## RESULTS

### (a) Externally driven jellyfish can double swimming speeds *in situ*

From plots of the vertical displacement over time (see Fig. 4 for a representative plot, with additional plots available in Figs. S2 and S3 in the Supplemental Material), we calculated vertical swimming speeds for three swim controller frequencies: 0 (control with inactive electrodes), 0.50, and 0.75 Hz for *N =* 2 animals, plotted in blue and red, as shown in Figure 6A. Because native animal pulses were not arrested using physical ablation or chemicals to reduce the animals’ biological pacemaker activity, Figure 6B shows the same vertical swimming speeds plotted over the measured swimming frequency (which illustrates the summative effect of the externally driven swim controller frequency and native animal pulses). The maximum vertical swimming speed obtained was 6.6 ± 0.3 cm s^-1^, externally driven at 0.75 Hz, compared to the minimum speed of 2.1 ± 0.1 cm s^-1^ in the absence of external frequency stimulation. Both the maximum and minimum speeds were observed in the same animal (animal 1, labeled in blue), which had a bell diameter of 11.3 ± 1.4 cm with a fineness ratio (defined as the ratio of the bell height to the bell diameter) of 0.39. See Table S1 in the Supplemental Material for more information on the experimental parameters and tabular results.

**Figure 6.**
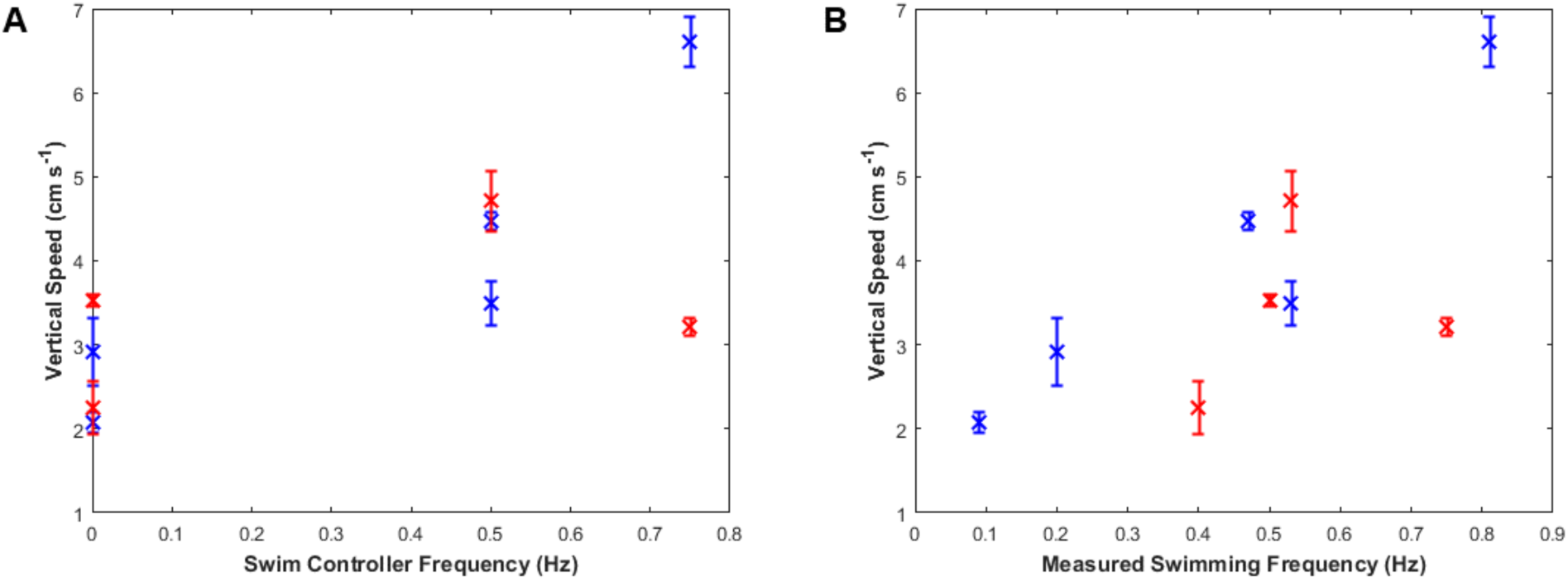
Vertical swimming speeds. (A) Vertical swimming speeds over swim controller frequency, the externally driven frequency set by the robotic system. Each animal is represented by a different color (blue or red). Two controls measurements were taken for each animal (at 0 Hz), and two videos were recorded at 0.50 Hz for animal 1 (blue). (B) Vertical swimming speeds over measured swimming frequency, the summation of both the externally driven frequency set by the robotic system and the animals’ own native pulses. Each animal is represented by a different color (blue or red). Variations in the animals’ baseline frequency can determine limits for robotic manipulation.

Swimming speeds generally increased with increasing frequency, although higher frequencies can decrease swimming speeds, as shown by the red data point at an externally driven frequency of 0.75 Hz in Figure 6A. This result confirms previous results of vertical swimming experiments in the laboratory, which showed peak swimming speeds at swim controller frequencies of 0.50 or 0.62 Hz [32]. To compare, the enhancement factors (the swimming speed divided by a baseline swimming speed in which the microelectronic system is embedded but inactive, i.e., the control case at 0 Hz) of both field data and prior work in the laboratory are plotted in Figure 7.

**Figure 7.**
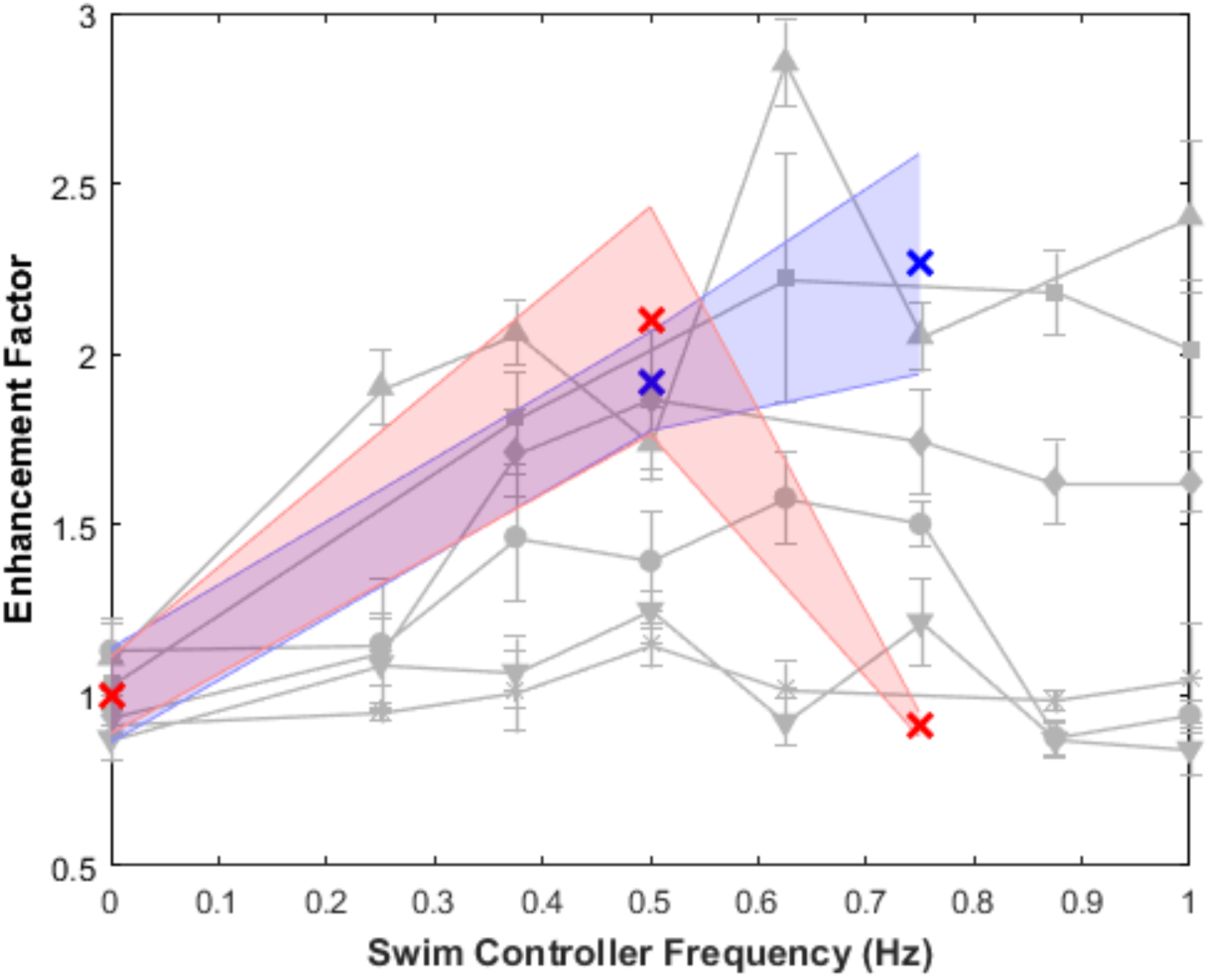
Enhancement factors measured in field work compared to prior work in the lab. The enhancement factor is defined as the swimming speed of each trial divided by the control case at 0 Hz, in which the swim controller is embedded but inactive. Experimentally driven frequency trials for each animal (*N =* 2, shown in blue or red) has been normalized to its own control trial. Prior laboratory work from similar vertical swimming experiments is shown in gray as a comparison, with symbol shape representing each individual animal (*N =* 6) [32]. Variability in the enhancement factor is influenced by the animals’ baseline swimming frequency, in the absence of stimulation.

The blue data point at 0.75 Hz shows the highest recorded swimming speed of the dataset, with a linear trend in vertical speed as measured swimming frequency increased. Because this biohybrid robotic jellyfish was not tested at higher external frequencies, such as 0.88 or 1.00 Hz, it is unclear whether the maximum enhancement occurred at 0.75 Hz or at which frequency the speed would maximize otherwise. Kinematic analyses of the bell morphology over contraction and relaxation times suggest that a maximum would occur no greater than 1.4 Hz, a proposed biological limit from previous research on muscle refractory periods in scyphozoan physiology [37]. However, at unusually high frequencies driven by the swim controller, such as 1.00 Hz, the bell morphology shifts to a more contracted phase over a longer period of time, in which the muscle ring cannot relax before the subsequent contraction, thereby decreasing the subumbrellar volume to decrease thrust and swimming speeds [32]. This bell morphological change was clearly observed in one animal, externally driven at 1.00 Hz, from the second dive. In that experimental condition, the biohybrid robotic jellyfish had a visibly slower swimming speed and never traversed the entire depth to the ocean surface, as opposed to other trials.

The effects of background flow on animal displacements were also calculated, with an example illustrated in Figure 5B to show oscillatory horizontal displacements resulting from surface currents (with wind speeds of 3.9 m s^-1^, West-southwest), compared to non-oscillatory vertical displacements (example in Fig. 4). As shown in Figure 5A, the main component of animal displacement was in the vertical direction.

In addition to the swimming performance of the overall biohybrid system, the microelectronic components were capable of performing for over 1.5 hours when run at 0.50 and 0.75 Hz, and over 45 min when run at 1.00 Hz, entirely submerged and exposed to natural conditions. Furthermore, the microelectronic system stayed embedded in the animals during each set of experiments despite physical handling and flow conditions (for a total of 15 min per system) until user removal for subsequent tests.

### (b) Comparison of theoretical and experimental swimming speeds

To determine whether theoretical models can predict swimming speeds for future applications to improve user controllability of the system, hydrodynamic models were run using input parameters from the videos of the highest measured swimming speeds for each animal, and run at all measured frequencies of that animal. As shown in Figure 8, the theoretical swimming speeds matched the trends in experimental swimming speeds, with mean differences between theoretical and mean experimental vertical speeds of 1.0 ±1.2 cm s^-1^ and 0.7 ± 0.6 cm s^-1^ for each animal, respectively. In addition to capturing the trends in swimming speeds, the model predicts the variations in swimming performance between the two animals at 0.75 Hz, including greater sensitivity to frequency changes in animal 1, as opposed to decreased sensitivity in animal 2.

**Figure 8.**
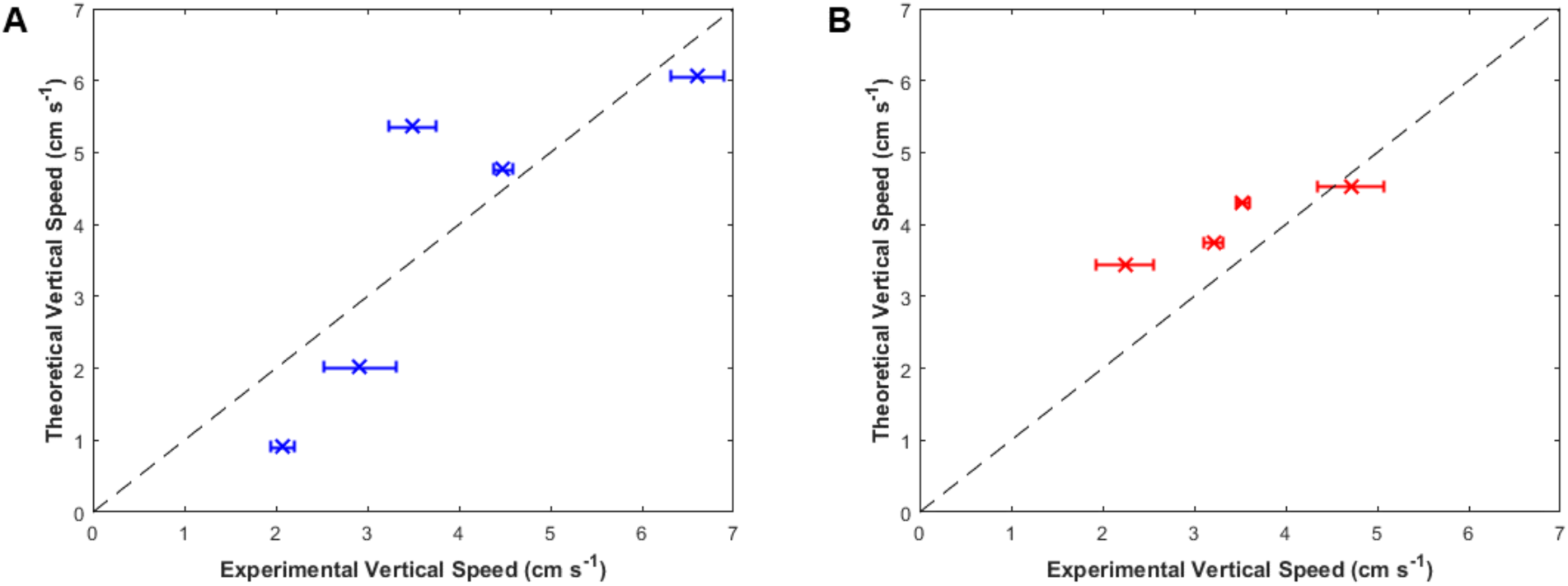
Theoretical versus experimental vertical swimming speeds. Theoretical vertical speeds from the hydrodynamic model versus the measured experimental vertical speeds for each individual jellyfish (A) blue using the input parameters delineated in the top row of Table 2 and (B) red using the input parameters delineated in the bottom row of Table 2. Inputs were obtained using morphological and time-dependent parameters for each jellyfish. The line of unity is plotted as black dashed lines.

## 4. DISCUSSION

The results of this *in situ* study suggest that biohybrid robotic jellyfish exhibit enhanced swimming modes, even in the presence of real-world conditions. Maximum enhancement factors for the *N =* 2 animals in field experiments were 2.3 ± 0.3 and 2.1 ± 0.3, and absolute swimming speeds increased two- to threefold. Despite limited field data, this corroborates laboratory experiments that reported user control of jellyfish swimming frequencies to enhance swimming speeds over twofold. Furthermore, comparable swimming speed enhancements in the field show a proof of concept that we can predictably improve jellyfish swimming speeds, even with background flows caused by winds of 3.9-4.6 m s^-1^ that resulted in oscillatory horizontal motion (Fig. 5), and potential interactions with other animals, such as the fish, ctenophores, and other medusae present in field experiments (Fig. 3).

This study also shows the robustness and reliability of the robotic system in these real-world conditions. The electronics for all four animals were viable for at least 45 min to 1.5 hours submerged in natural saltwater, dependent on the stimulation frequency. Future studies can conduct field experiments in more locations, including in open water or at greater depths farther from the shore.

To create a more user-controllable biohybrid robotic system, we need to comprehensively study both the natural animal system and how the robotic system interacts with the animal. For example, endogenous swimming frequencies occurred from 0.09 to 0.50 Hz in the absence of external frequency control. However, this range is narrower when we consider each animal separately; the natural pulse response observed in one individual animal ranged from 0.09 to 0.20 Hz, whereas the response in a second animal ranged from 0.40 to 0.50 Hz. Higher frequencies might have been the result of natural animal variations and increased sensory information from chemical or mechanical stimulation in the ocean. The differences in the animals’ baseline swimming frequencies in Figure 6 suggest that animals with lower natural pulses are more sensitive to robotic stimulation, which suggests that future studies can use animals that naturally exhibit lower swimming frequencies to maximize enhancement. This could also explain the variations in enhancement factors among the laboratory results in prior work [32].

Furthermore, the animal with a smaller native frequency exhibited greater absolute swimming speeds, as plotted in Figure 6, with comparable enhancement factors to prior animals (in gray) in Figure 7. Animal 2 (red) had a larger fineness ratio of 0.43 than animal 1 (blue), which had 0.39. However, the speed enhancements of the present study did not surpass prior enhancements. This suggests that in addition to parameters such as size and fineness ratio [32,38,39], other morphological and time-dependent parameters can be critical when choosing individual animals for future work.

The hydrodynamic model we describe can be used to determine which animals are appropriate for optimal robotic integration. By using morphological and time-dependent input parameters from the videos of these animals at only one swimming speed, we predicted the vertical swimming speeds at all frequencies, with a mean error of 0.8 cm s^-1^ (Fig. 8). The model captures animal behavior at each of the nine test cases, including predicting doubled enhancements at 0.50 Hz for both animals, as well as the disparity at 0.75 Hz between increased speed in animal 1 and decreased speed in animal 2. Although this jetting model does not incorporate the full hydrodynamics of rowing propulsion evident in *A. aurita*, this simple model is a useful first order prediction. Because the results of these experiments validate the hydrodynamic model and trends in swimming speeds, further studies can systematically determine which bell morphological parameters most affect swimming speeds or other metrics of maneuverability through both theoretical modeling and experiments. Regardless, the current model shows utility by predicting the swimming speeds and variations in both animals.

Regarding maneuverability, the current study is limited to purely vertical swimming, ballasted by the swim controller to maintain its upright position. However, future studies can use an unstably balanced weighting system and asymmetric activation of electrodes to allow turning. Accelerometers on both the animal and camera systems can also be used to track 3D motion of the biohybrid robotic jellyfish for more complicated jellyfish maneuvering, such as following trajectories with closed-loop controls.

The present study also examined horizontal swimming speeds as a proxy for background flow conditions, by taking advantage of coastal conditions to assume primarily horizontal surface currents [40]. These horizonal ocean currents were less likely to affect the vertical swimming speeds exhibited by the biohybrid robotic jellyfish with their ballasted design. Additionally, experimental trials were conducted successively in a narrow span of time to minimize more extreme variations in flow conditions among subsequent trials, with both dives occurring for one hour per day. Future *in situ* studies can determine how various background flows affect jellyfish swimming using particle image velocimetry, and more laboratory experiments to systematically characterize the user control of jellyfish swimming can also include studies of controlled background flows and their effects on swimming speeds.

Finally, the main limitation of the current work is the small sample size due to challenges in field work and conditions. Nevertheless, the results demonstrate a proof of concept that a biohybrid robotic jellyfish system can perform at doubled speeds predictably *in situ*, with the potential for wider use in ocean monitoring after further design modifications. User control of jellyfish swimming has been established for unidirectional swimming in prior and current work. By using the biohybrid robotic jellyfish system in this work as a basis, future experiments can focus on animal maneuverability and robotic design. Suggestions include determining the electrode stimulation patterns needed for asymmetrical swimming and trajectory tracking in the laboratory, adding sensors to collect data from the environment, and integrating biodegradable electronic components for field measurements.

## 5. CONCLUSIONS

The present study demonstrates a proof of concept that biohybrid robotic jellyfish can be implemented in coastal conditions, with doubled swimming speed enhancements, comparable to prior experiments conducted in the laboratory. Differences in the animals’ baseline swimming frequencies could determine sensitivity to robotic manipulation, to address the variability seen in both current and prior work. A theoretical model was developed to predict experimental swimming speeds with mean errors of 0.8 cm s^-1^, using input parameters estimated from videos of one trial to extrapolate speeds at all frequencies for that individual animal. The model accurately predicted variability in swimming speeds among the animals to provide a basis for choosing which animals would be optimal for robotic manipulation in the future. Therefore, this work addresses open questions in the user control of jellyfish swimming, including how real-world environments affect swimming speed enhancements observed in the laboratory, which factors cause large animal variability, and whether theoretical models can predict which individual animals perform better.

Because the biohybrid robotic jellyfish in this study have operated with predictable user control under field conditions, future work can use this existing microelectronic and live animal system *in situ* as an alternative method to monitor the ocean. By improving maneuverability and incorporating sensors to track environmental changes (such as salinity, acidity, and temperature) into the present design, we can potentially use biohybrid robotic jellyfish as a ubiquitous and energy-efficient tool.

## DATA AVAILABILITY

Data are available in the Stanford Digital Repository: https://purl.stanford.edu/mh950tt5866

## ACKNOWLEDGMENTS

We would like to acknowledge Cabrillo Marine Aquarium for providing *A. aurita* medusae, and Valerie A. Troutman and Jennifer L. Cardona for their help in shipping animals from Stanford, CA to Woods Hole, MA.

## AUTHOR CONTRIBUTIONS

NWX and JOD conceived the study and edited the manuscript; BG and NWX conducted preliminary tests in the Atlantic Ocean to preempt diver field experiments; JPT, JJC, and SPC conducted subsequent field experiments as scientific scuba divers; NWX conducted field experiments from the laboratory and on shore, performed the data analysis, and wrote the initial manuscript.

