## Supplemental Material for "Field testing of biohybrid robotic jellyfish to demonstrate enhanced swimming speeds"

**Table S1. Experimental parameters and results for field experiments.** The diameter, fineness ratio (defined as the ratio of the bell height to diameter), swim controller frequency, measured frequency from counting animal pulses, and calculated vertical swimming speeds for $N = 2$ animals (labeled as blue and red to match plot figures).

| *Animal* | *Diameter (cm)* | *Fineness Ratio* | *Swim Controller Frequency (Hz)* | *Measured Frequency (Hz)* | *Vertical Swimming Speed (cm/s)* |
|---|---|---|---|---|---|
| 1 (blue) | 11.3 ± 1.4 | 0.39 | 0 (control) | 0.09 | 2.1 ± 0.1 |
| 1 (blue) | 11.3 ± 1.4 | 0.39 | 0.50 | 0.53 | 3.5 ± 0.3 |
| 1 (blue) | 11.3 ± 1.4 | 0.39 | 0.50 | 0.47 | 4.5 ± 0.1 |
| 1 (blue) | 11.3 ± 1.4 | 0.39 | 0 (control) | 0.20 | 2.9 ± 0.4 |
| 1 (blue) | 11.3 ± 1.4 | 0.39 | 0.75 | 0.81 | 6.6 ± 0.3 |
| 2 (red) | 9.8 ± 0.8 | 0.43 | 0 (control) | 0.40 | 2.2 ± 0.3 |
| 2 (red) | 9.8 ± 0.8 | 0.43 | 0.50 | 0.53 | 4.7 ± 0.4 |
| 2 (red) | 9.8 ± 0.8 | 0.43 | 0 (control) | 0.50 | 3.5 ± 0.1 |
| 2 (red) | 9.8 ± 0.8 | 0.43 | 0.75 | 0.75 | 3.2 ± 0.1 |

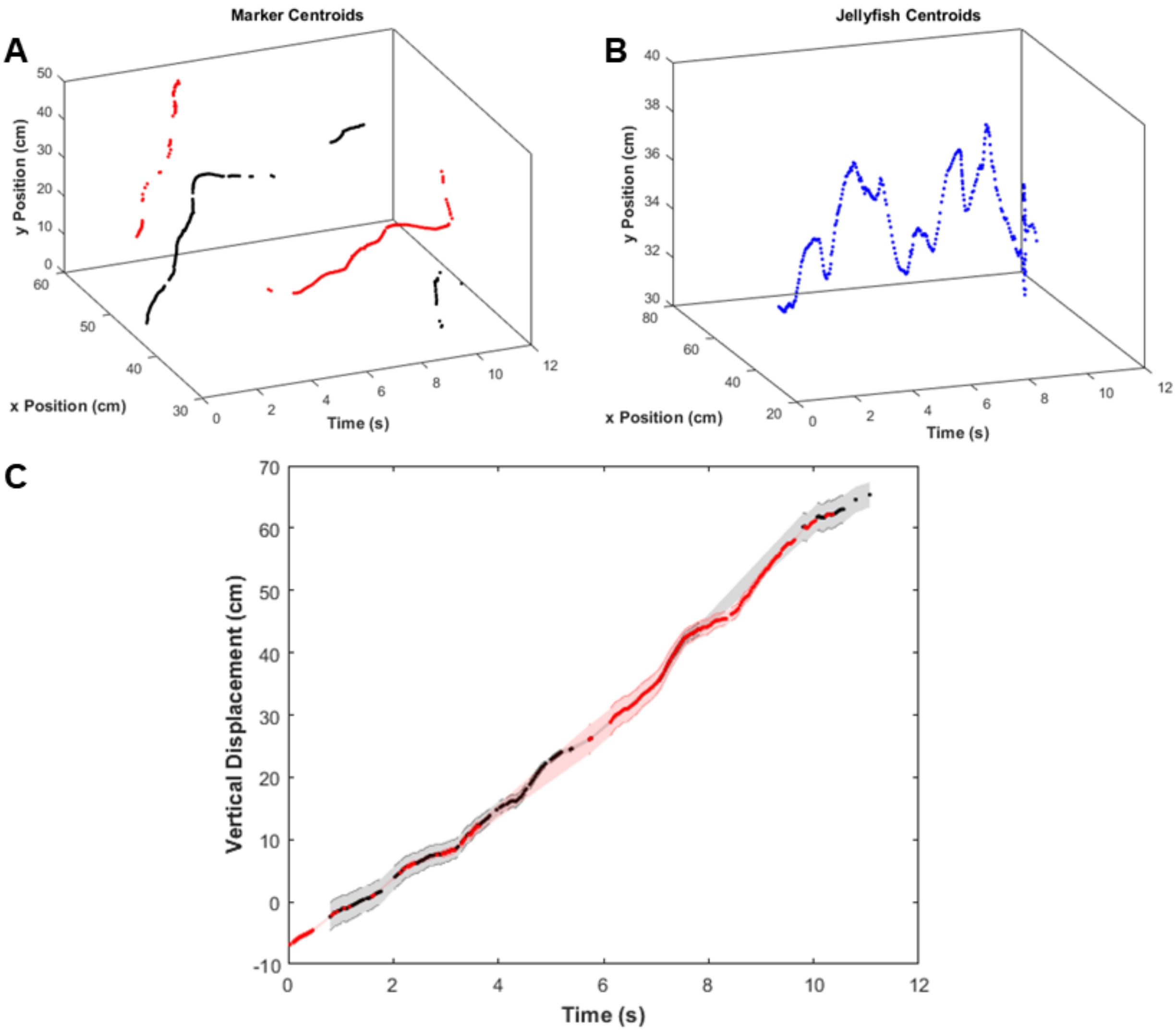


**Figure S1. Representative plots tracking animal displacements over time to calculate vertical swimming speeds.** To calculate the animals' displacement over time with respect to the rope, a prominent background feature as a ground truth, we first tracked (A) centroids of the red and yellow markers on the rope in each image frame. A sample from one video (animal 1 driven at 0.75 Hz) is shown, with red markers plotted in red and yellow markers plotted in black for improved visualization. (B) Centroids of the jellyfish (by tracking the blue polypropylene housing) in each image frame over time. (C) Vertical displacement over time of the jellyfish with respect to the rope (data also presented in Figure 4 of the main), with the error propagated from conversions in pixel space to centimeter space. Tracks were assembled by stitching vertical positions using both red and yellow markers (shown in red and black, respectively), to show accuracy in overlap.

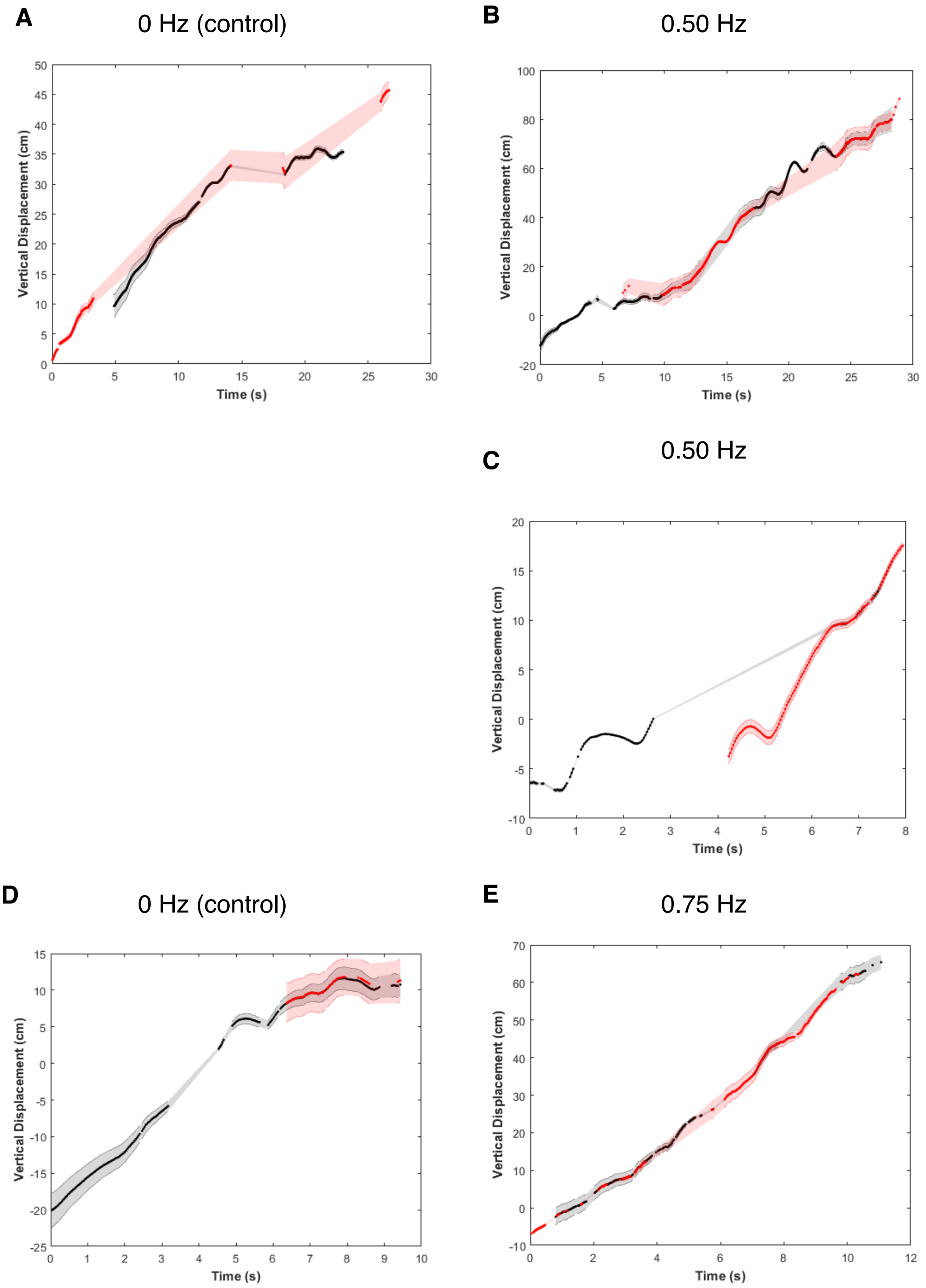
A
0 Hz (control)
B
0.50 Hz
C
0.50 Hz
D
0 Hz (control)
E
0.75 Hz
Vertical Displacement (cm)
Time (s)

**Figure S2. Displacements over time for jellyfish, animal 1.** Vertical displacement over time of the jellyfish with respect to the rope, with the error propagated from conversions in pixel space to centimeter space, assuming pixel-level accuracy in centroids. Tracks were assembled by stitching vertical positions using both red and yellow markers (shown in red and black, respectively), to show accuracy in overlap.

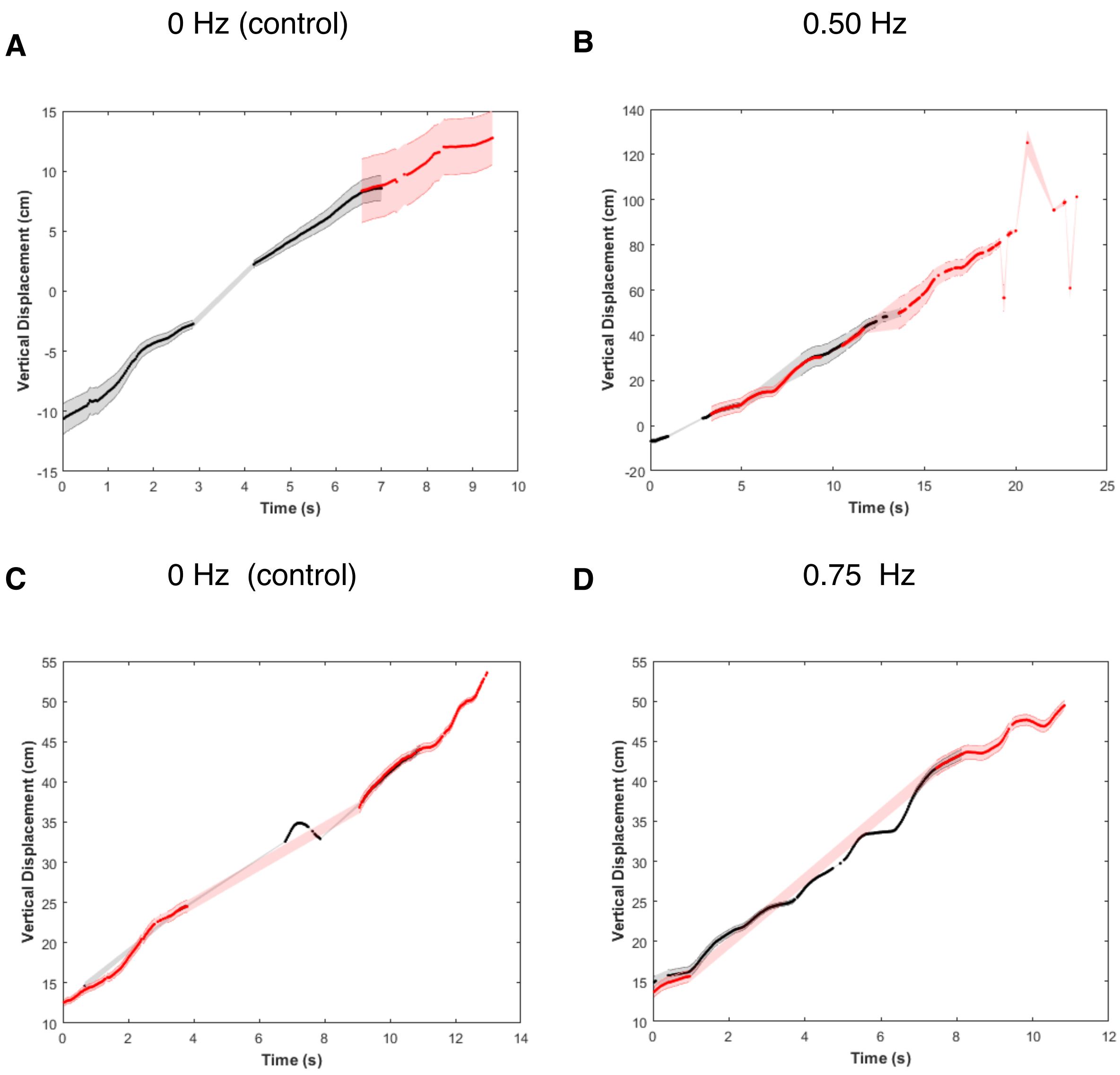


**Figure S3. Displacements over time for jellyfish, animal 2.** Vertical displacement over time of the jellyfish with respect to the rope, with the error propagated from conversions in pixel space to centimeter space, assuming pixel-level accuracy in centroids. Tracks were assembled by stitching vertical positions using both red and yellow markers (shown in red and black, respectively), to show accuracy in overlap.

**Ethical considerations**

Although jellyfish are invertebrates that are not under consideration of the Institutional Animal Care and Use Committee (IACUC), the authors would like to address ethical concerns about these animal experiments and incorporating live animals into biohybrid robotic constructs.

Jellyfish do not have a centralized nervous system and have no known pain receptors. There is no research that suggests that these animals can feel pain, but regardless, we know that jellyfish secrete excess mucus when stressed. If we use their stress response as a proxy for pain, animals did not exhibit stressed behavior during or after experiments. Furthermore, animals did not show any side effects after the robotic devices were removed; animal behavior returned to its normal state, including normal feeding behaviors.

Regarding introducing more animals into different ecosystems in the ocean, *A. aurita* are naturally found in Woods Hole. However, because no animals were observed during field experiments, we introduced *A. aurita* and monitored the biohybrid robotic jellyfish carefully to ensure no jellyfish or electronic debris were left in the ocean after experiments. To address these issues in future experiments, further work can include using jellyfish found in their natural habitats, given the ubiquity of *A. aurita*, and incorporating biodegradable electronics already used for medical purposes.
